# A semi-automated method for quantifying optokinetic reflex tracking acuity

**DOI:** 10.1101/2023.08.03.551461

**Authors:** James K. Kiraly, Scott C. Harris, Timour Al-Khindi, Felice A. Dunn, Alex Kolodkin

**Affiliations:** Solomon H. Snyder Department of Neuroscience, The Johns Hopkins Kavli Neuroscience Discovery Institute, The Johns Hopkins University School of Medicine; Department of Ophthalmology, University of California, San Francisco

## Abstract

The study of murine behavioral responses to visual stimuli is a key component of understanding mammalian visual circuitry. One notable response is the optokinetic reflex (OKR), a highly conserved innate behavior necessary for image stabilization on the retina. The OKR provides a robust readout of image tracking ability and has been extensively studied to understand the logic of visual system circuitry and function in mice from different genetic backgrounds. The OKR consists of two phases: a slow tracking phase as the eye follows a stimulus to the edge of the visual plane, and a compensatory fast phase saccade that maintains the image within the visual field. Assessment of the OKR has previously relied on counting individual compensatory eye saccades to estimate tracking speed. To obtain a more direct quantification of tracking ability, we have developed a novel, semi-automated analysis program that allows for rapid and reproducible quantification of unidirectional tracking gains, in addition to being adaptable to any video-oculography equipment. Our analysis program allows for the selection of slow tracking phases, modeling of the vertical and horizontal eye vectors, quantification of eye movement relative to the stimulus, and organization of resultant data into a usable spreadsheet for statistical and graphical comparisons. This quantitative and streamlined analysis pipeline provides a faster and more direct measurement of OKR responses, thereby facilitating further study of visual behavior responses.

**SUMMARY:** We describe here a semi-automated quantitative analysis method that directly measures eye tracking resulting from murine visual system responses to two-dimensional image motion. A Python-based user interface and analysis algorithm allows for higher throughput and more quantitative measurements of eye tracking parameters than previous methods.

## INTRODUCTION

Image stabilization relies on precise oculomotor responses to stabilize a moving object on the visual field. This stabilization is driven primarily by two motor responses: the optokinetic reflex (OKR) and the vestibulo-ocular reflex (VOR)^1-3^. Slow object motion across the retina induces the OKR, which elicits reflexive eye rotation in the corresponding direction to maintain the image within the visual field^1,2^. This movement, known as the slow phase, is interrupted by compensatory saccades, known as the fast phase, in which the eye rapidly resets in the opposite direction to allow for a new slow phase. Whereas the VOR relies on the vestibular system to elicit compensatory head movements^3^, the OKR is initiated in the retina by directional firing of direction-selective retinal ganglion cells (DSGCs) and subsequent signaling to the Accessory Optic System (AOS) in the midbrain^4,5^. Due to its direct reliance on retinal circuits, the OKR has been frequently used to determine visual tracking ability in both research and clinical settings^6,7^.

The OKR has been studied extensively as a tool for assessing basic visual ability^2,6,8^, DSGC development^9-12^, oculomotor responses^13^, and physiological differences between genetic backgrounds^7^. The OKR is evaluated in head-fixed animals to which a moving stimulus is presented^14^. Oculomotor responses are captured using an infrared camera and eye tracking motions are captured as OKR waveforms in horizontal and vertical directions^9^. Previous studies primarily quantify OKR strength based on the frequency of fast phase saccadic motions since increased saccade number correlates with increased slow phase tracking speeds^7^. Although this provides a fairly accurate estimation of tracking ability, this method relies on an indirect metric to quantify the slow phase response and introduces a number of biases including an observer bias in saccade determination, a reliance on temporally consistent saccadic responses across a set epoch, and an inability to assess the magnitude of the slow phase response.

In order to address these concerns with current OKR assessment approaches and to satisfy the need for an in-depth quantification approach for OKR parameter assessment, we have developed a new analysis method to quantify OKR waveforms that uses high throughput and accessible Python-based software. Through a semi-automated analysis routine, modeling and quantification of OKR slow phase responses can be studied in greater depth, allowing for automated calculation of eye velocity and gains to provide a rigorous and reproducible direct measurement of tracking ability.

### PROTOCOL

All animal experiments performed at The Johns Hopkins University School of Medicine were approved by the Institutional Animal Care and Use Committee (IACUC) at The Johns Hopkins University School of Medicine (JHUSOM). All experiments performed at the University of California, San Francisco (UCSF) were performed in accordance with protocols approved by the University of California, San Francisco Institutional Animal Care and Use Program.

#### Behavioral Data Collection

1. Record OKR wave data using video-oculography method of choice.

Representative data collected at JHUSOM were obtained using the headpost implantation surgery and video-oculography method as described^9,13^ (**Figure 1a**). Representative data collected from UCSF were obtained through the video-oculography method as described in Harris and Dunn^10^ (**Figure 5a**)
1.1 Make note of stimulus and recording parameters: the speed and length of the stimulus and the frame rate of the recording camera to ensure accuracy of later gain calculations.
2. Export wave data as a .CSV file containing horizontal and vertical wave data.

Representative data from JHUSOM was processed via Igor Pro v6.37 and exported as .CSV files using a custom script.
2.1 Wave data should be organized as a tab-delimited CSV file with two columns containing horizontal data (“epxWave”) and vertical data (“epyWave”)

**Figure 1.**
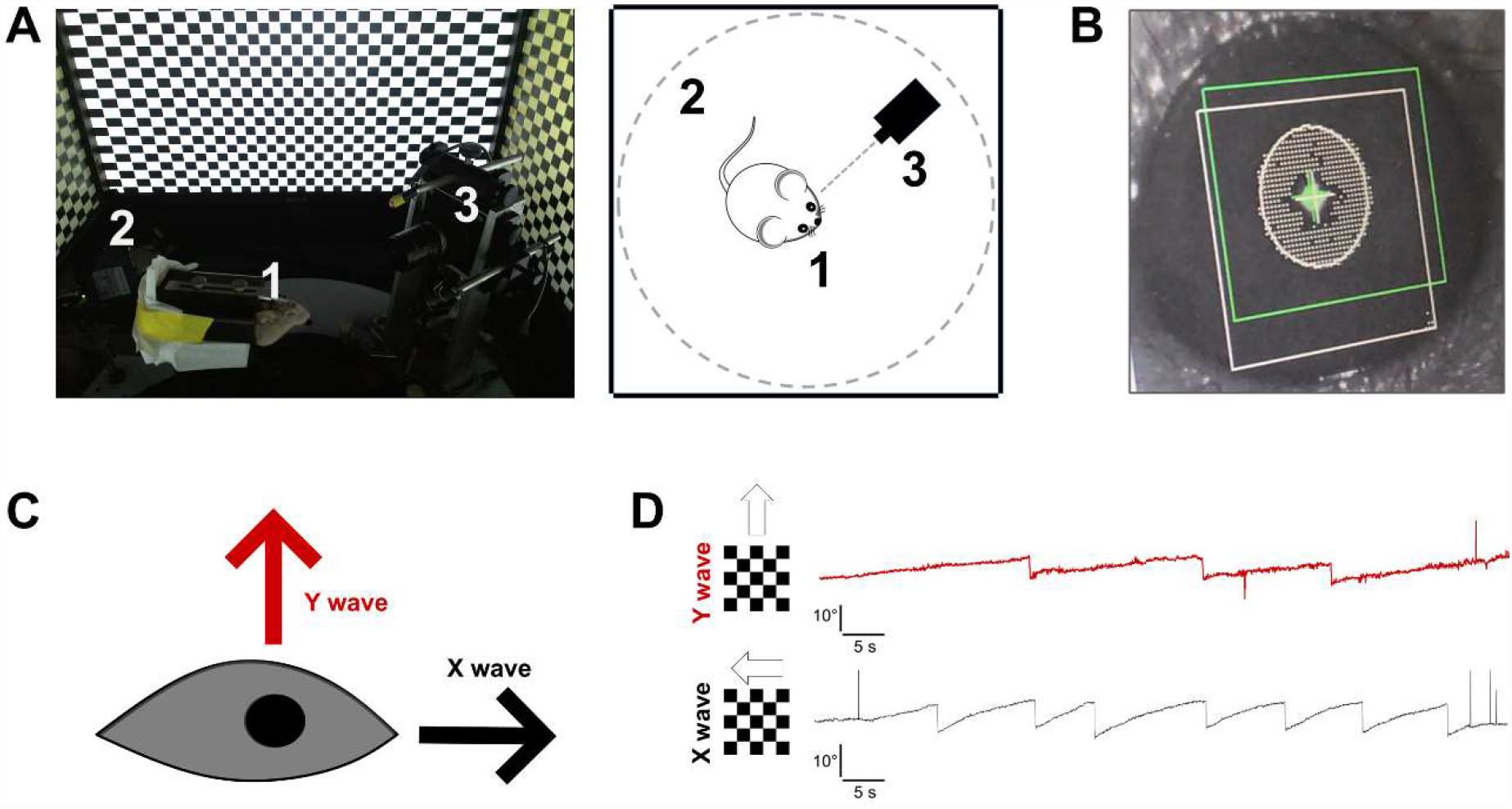
Collection of OKR response data. **A)** Set-up of OKR virtual drum for behavioral stimulation as described^9,13^. Four monitors surround a head-fixed animal (1), displaying a continuously moving checkerboard stimulus (2). The virtual drum can present unidirectional movement in all four cardinal directions as well as oscillatory sinusoidal stimuli. The mouse’s left is illuminated by an infrared (IR) light and recorded with a camera (3) to record visual system responses reflected in eye tracking. **B)** Analysis of eye tracking occurs through capturing of the pupil and a corneal reflection generated by the IR light. Data collection and calculation of eye movements in response to the virtual drum were performed as described in ^9,13^. **C)** Schematic of eye vectors moving vertically (Y wave) and horizontally (X wave). **D)** Sample traces of an eye’s tracking response to upward motion and to backwards motion.

#### Installation of analysis software

1. Download and install Python (https://www.python.org/) and Spyder (https://www.spyder-ide.org/)

1.1 Installation of Python and Spyder via Anaconda is recommended for interactive supervision of waves.
2. Create a virtual environment in Python (NOTE: Python versions ≥ 3.8.3 are recommended)

2.1 Install required packages:

- PyQT5
-Scipy
-Matplotlib
-Sklearn.neighbors
-Scikit-learn
-Pandasgui
3. Download the newest version of the Python script (e.g. *PyOKR_v2*.*3_win*.*py*) according to the OS you are using from the Kolodkin laboratory GitHub (https://github.com/KolodkinLab/PyOKR).

NOTE: sample data can also be downloaded in this GitHub under “Sample Data”
4. Open the script within Spyder and connect it to your virtual environment.

#### Analysis of wave data

To verify successful installation of the software, use the provided sample data for analysis. Below, we describe an analysis tutorial using sample wave data. An in-depth tutorial can be found on the Kolodkin laboratory’s GitHub (https://github.com/KolodkinLab/PyOKR).

**1. Initialization of analysis and file imports**

1.1 Run the entire script and a user interface will appear for wave analysis.
1.2 Using the button **Set CSV**, or the command **Ctrl+O**, will result in a browser appearing that will allow the user to select the desired wave file.
1.3 Using the button **Set Output folder**, or the command **Ctrl+E**, a folder browser will appear allowing for selection of the output file. This will be the folder into which the final analysis for a given mouse will be exported.
1.4 Input the final analysis file name under **Output file**. We recommend a format such as: *MouseGenotype_MouseNumber_Analysis*.
1.5 Set the program for an individual mouse using the command **Set Mouse** under **File**, or command **Ctrl+S**. This initializes the dataset containing the data for an individual mouse and should be performed only once per mouse.
**2. Definition of wave file parameters**

Parameters for wave analysis must be defined based on the presented stimulus for a given wave file. Directional parameters are first set to represent the four cardinal directions.
Stimulus speed in degrees per second are also set manually based on the collection method employed. Epoch time intervals for active stimuli must be adjusted within the code according to the collection method. Default values are set to five 30-second epochs and a stimulus speed of five degrees per second.
2.1 Directionality is defined under **Select stimulus direction** as either a **Horizontal** or a **Vertical** stimulus.
2.2 Rotational direction is defined under **Select stimulus rotation** as either clockwise (**CW**) or counterclockwise (**CCW**). For horizontal stimuli, **CW** refers to temporal-to-nasal motion and **CCW** refers to nasal-to-temporal motion. For vertical stimuli, **CW** refers to downward motion and **CCW** refers to upward motion.
NOTE: the description of stimulus directions can be modified in the code to fit specific collection methods.
2.3 A given epoch for a particular stimulus is set under **Select epoch**. Based on intervals of presented stimuli, this allows the user to scan through regions of presented stimulus and record tracking gains.
2.4 Stimulus speed is set under **Enter stimulus speed** as a numerical value in degrees per second.
**3. Supervised selection of tracking phases**

3.1 To identify regions of slow tracking, fast phase saccades are automatically selected under **Preliminary ETM plot**. The tracking wave should appear in red with blue dots automatically placed on saccadic movements. Saccades are identified through rapid changes of slope values and a kernel density estimation to determine maximal points of change. Due to variability in wave quality, manual supervision can be used to adjust these automatically placed points marking saccades. **Left Mouse Button** removes points (false positives) and **Right Mouse Button** adds points (false negatives). When fast phase saccades are adequately selected, **Middle Mouse Button** will save points and the graph can be closed.
NOTE: Use of a computer mouse is recommended for fast supervision.
3.2 Boundaries of the saccade are then set using **Top point adjustment** and **Bottom point adjustment**. Top and bottom points of a saccade are found by scanning backward and forward from the preliminary saccade points. Wave inflection points are calculated and the top and bottom points are automatically placed. Adjustments can be made with the same control scheme described above using **Preliminary ETM Plot**. When points are properly placed, use the **Middle Mouse Button** to save points.
**4. Analysis of slow tracking phases**

After slow phases are appropriately selected as described, analysis of the waves is performed automatically. Waves are analyzed through a polynomial approximation of selected slow phases using an arclength of the polynomial to calculate the distance traveled in XY directions. Distances traveled over time are used to define the velocity, and tracking gains are found based on the velocity relative to the stimulus speed. Calculations are conducted as a vector sum of XY motion and are also broken down into horizontal and vertical vector components.
4.1 Polynomial order can be defined under **Set Polynomial Order** based on desired fitting of the polynomial to the wave. For tracking gains, a single order polynomial is recommended to accurately represent eye velocity relative to the linear speed of the stimulus. However, higher order polynomials can be used for a more accurate representation of eye movement velocity.
4.2 Once parameters are properly set, **Analyze current ETMs** will display the final selection of slow phases (**Figure 2a-d**) and will calculate the distances, velocities, and tracking gains averaged across the epoch (**Figure 2e**). Visualization of the eye movement in 3D can be viewed using **3D graph** (**Figure 3b**).
4.3 Analysis of tracking data can be then added to an individual mouse dataset with **Add epoch**. Values from the given epoch will be stored based on the directional stimulus as defined earlier. Added values for a given mouse can be viewed with **View current dataset**. Averages are automatically calculated based off of stimulus grouping.
4.4 After an epoch is added, additional epochs for a given file can be selected and analyzed with **Select epoch**, following steps 3.1 to 4.3.
4.5 Once a wave file is fully analyzed, this process can be repeated by opening new files, setting appropriate parameters, and analyzing them accordingly. By repeating steps 2.1-4.3 for each file, a final dataset will be generated containing all wave data for a given mouse. It is essential to not reset the dataset with **Set Mouse** or to close the interface, which could reset the compiled dataset for that given mouse.
**5. Final export of data**

5.1 After data analysis is complete for a given mouse, with all directions analyzed, data can be exported via **Export data**. The raw dataset will be exported based on the **Output file** name set previously and saved along the path set by **Output Folder**.
5.2 As animals are analyzed the output folder will contain a number of CSVs containing tracking data. Data can be re-organized for easier analysis using the command **Sort Data** under the **Analysis** tab. This function will compile and sort all average values for all the analyzed animal file stored within the output folder to allow for easier generation of graphs and statistical comparisons.

**Figure 2.**
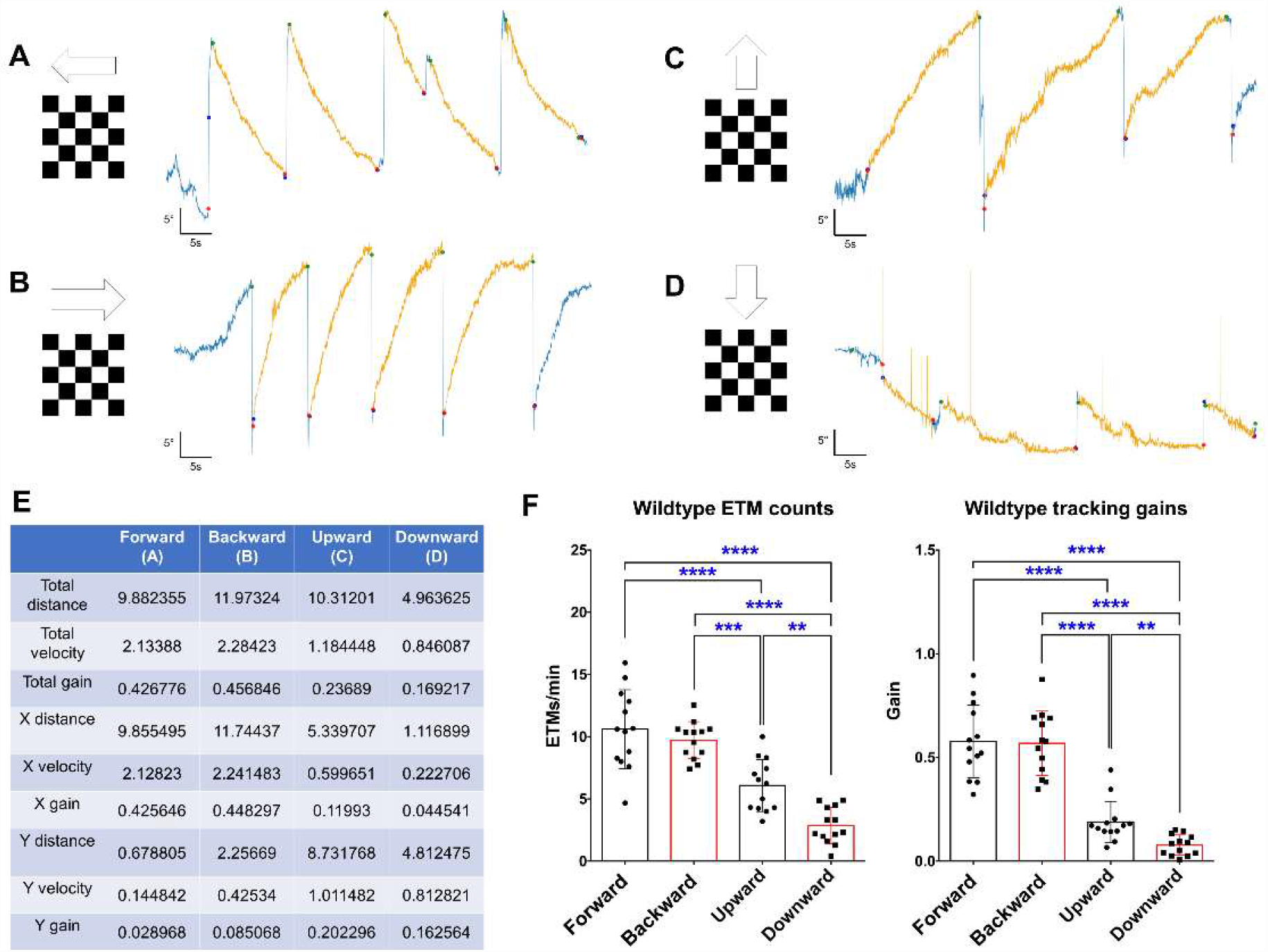
Tracking analysis of unidirectional visual responses. **A-D)** Identification and selection of slow tracking phases for gain analysis. Sample unidirectional traces are shown with visual responses to forward (**A**), backward (**B**), upward (**C**), and downward (**D**) motion. Slow phases are selected between fast phase saccades and highlighted in yellow. **E)** Quantification of previous traces (**A-D**). Average distance, velocity, and gain of slow phases are calculated for the total vector (XY) as well as broken down to horizontal (X) and vertical (Y) components. Distance calculated in degrees; velocity calculated in degrees/sec; gain calculated as velocity/stimulus speed. **F)** 3D projection of eye movement in horizontal and vertical space in response to upward stimulus (**C**). **G**) Calculated tracking gains of wildtype animals (n=14) in the four cardinal directions. Data are presented as mean ± SD. Data analyzed with a one-way ANOVA with multiple comparisons. *p<0.05, ****p<0.0001.

**Figure 3.**
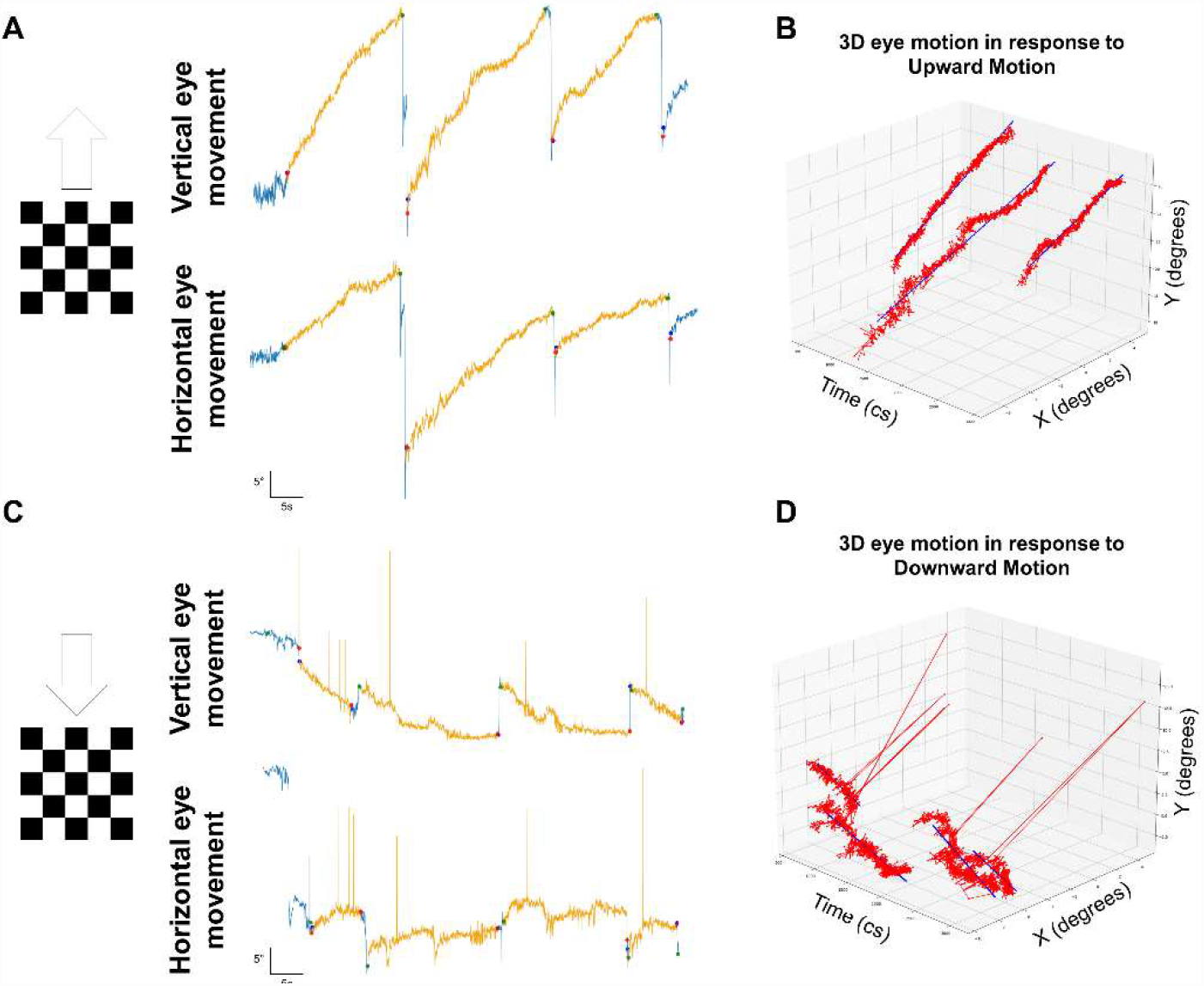
Directional tracking can be modeled in its horizontal and vertical components. **A)** Vertical and horizontal eye tracking waves in response to an upward stimulus. **B)** Three-dimensional model of the eye’s movement vector over time in response to upward motion. **C)** Vertical and horizontal eye tracking waves in response to a downward stimulus. **D)** Three-dimensional model of the eye’s movement vector over time in response to downward motion.

## REPRESENTATIVE RESULTS

To validate the analysis method described above, we quantified OKR tracking gain on wave traces collected from wild mice and a conditional knockout mutant with a known tracking deficit. In addition, to test our analysis method’s scalability, we analyzed wildtype traces derived using a different video-oculography collection method. Using recordings from unidirectional grating stimuli (**Figure 1d**), we calculated OKR tracking gains in the four cardinal directions (**Figure 2f**) for wildtype animals (n=17). Disparity in tracking ability relative to stimulus direction is consistently observed, with equally robust horizontal responses that demonstrate significantly higher tracking gains than vertical responses^2^. Further, asymmetric tracking gain between upward and downward responses are also observed, as has been previously reported^2,10^. The relative strength and consistency of tracking gains in comparison to published characterization of OKR responses indicate that tracking gains accurately reflect tracking ability. In addition to single directional gain calculations, horizontal and vertical eye movement can be modeled simultaneously (**Figure 3**), allowing for a three-dimensional reconstruction of eye movement in response to a given stimulus. This provides additional quantification capability that provides useful data for future studies that investigate cross-coupled horizontal and vertical responses.

To validate the use of this method in identifying significant behavioral changes between experimental conditions, we re-analyzed published data from Al-Khindi et al.^9^ to confirm that deficits in vertical tracking assessed in that study by manual counting of fast phase saccades can be reflected in tracking gains. Previous work shows that genetic inactivation in the retina of the transcription factor, *T-box Transcription Factor 5* (*Tbx5*) through conditional knockout using *Protocadherin 9-Cre* (*Pcdh9-Cre*) causes select loss of upward-tuned On Direction-Selective Ganglion Cells (up-oDSGCs), and that *Tbx5f/f; Pcdh9-Cre* mutants exhibit specific loss of vertical tracking^9^. Quantitative analysis using the method described here shows that *Tbx5* cKO animals retain normal horizontal tracking gains (**Figure 4a**), similar to those described previously obtained by manual counting of fast phase saccades (**Figure 2f**); however, these mice show significant loss of vertical tracking ability, with near zero gains in response to both upward and downward stimuli (**Figure 4b-c**). Recapitulation using our new methodology of the previously described phenotype demonstrates the precision and sensitivity of this method that allows for quantitative comparison of OKR responses in mice of different genetic strains.

**Figure 4.**
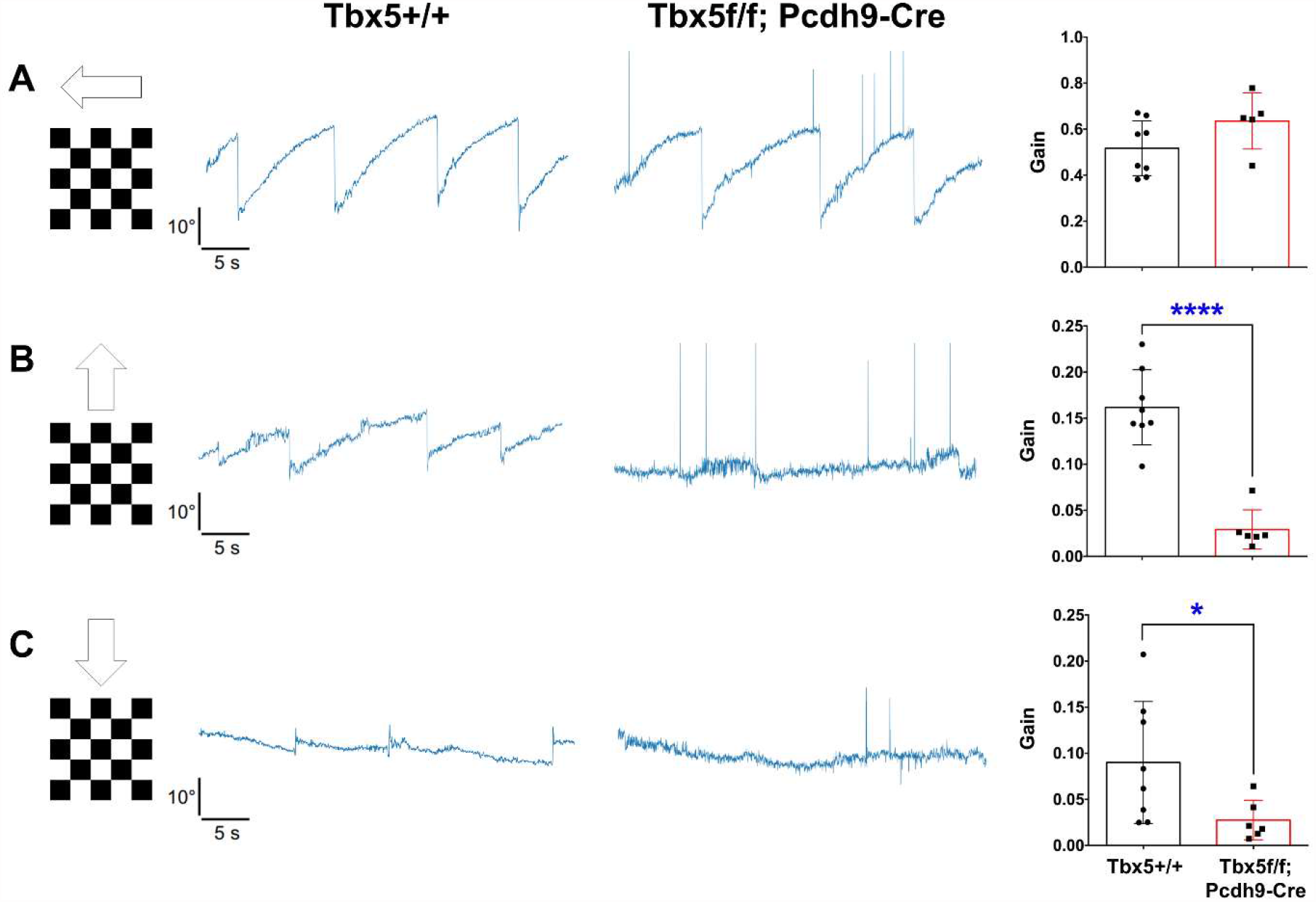
Assessment of *Tbx5f/f; Pcdh9-Cre* mice shows significant deficits in unidirectional vertical tracking gains. **A)** *Tbx5f/f; Pcdh9-Cre* animals show no significant change in horizontal tracking gain. **B-C)** *Tbx5f/f; Pcdh9-Cre* animals show a significant reduction of gain in their vertical responses. Data are presented as mean ± SD. Data analyzed with unpaired T-tests. *p<0.05, ****p<0.0001.

Finally, we analyzed wildtype vertical OKR traces collected at UCSF to validate our method’s utility with different video-oculography methods and stimulus parameters. Data from UCSF were collected using a hemispherical projection system in which moving bars are presented to the mouse through the reflection of a 405nm wavelength projector onto a hemisphere surrounding the fixed animal^10^ (**Figure 5a**). Unidirectional vertical gratings were presented at a speed of 10 degrees per second and mouse’s responses were recorded over 60 second intervals (**Figure 5b-c**). Following slight modifications to fit these different parameters of this experimental set-up, vertical traces were quantitatively analyzed (**Figure 5d**) and upward responses were compared to downward responses. Upward responses were significantly stronger than downward, as expected^10^; however, tracking gains were slightly reduced compared to traces recorded at JHUSOM (**Figure 2f**). This reduction can be attributed to differences between stimulus parameters, including increased stimulus speeds; however, the overall accuracy of this analysis indicates that our method is adaptable beyond our specific OKR data collection system and can be applied to any OKR recordings, independent of video-oculography methods. These results demonstrate that our new analysis method is accurate and can be generally applied to the study of oculomotor responses, allowing for precise quantitative comparisons among mice belonging to different groups to further the study of murine visual image stabilization circuitry.

**Figure 5.**
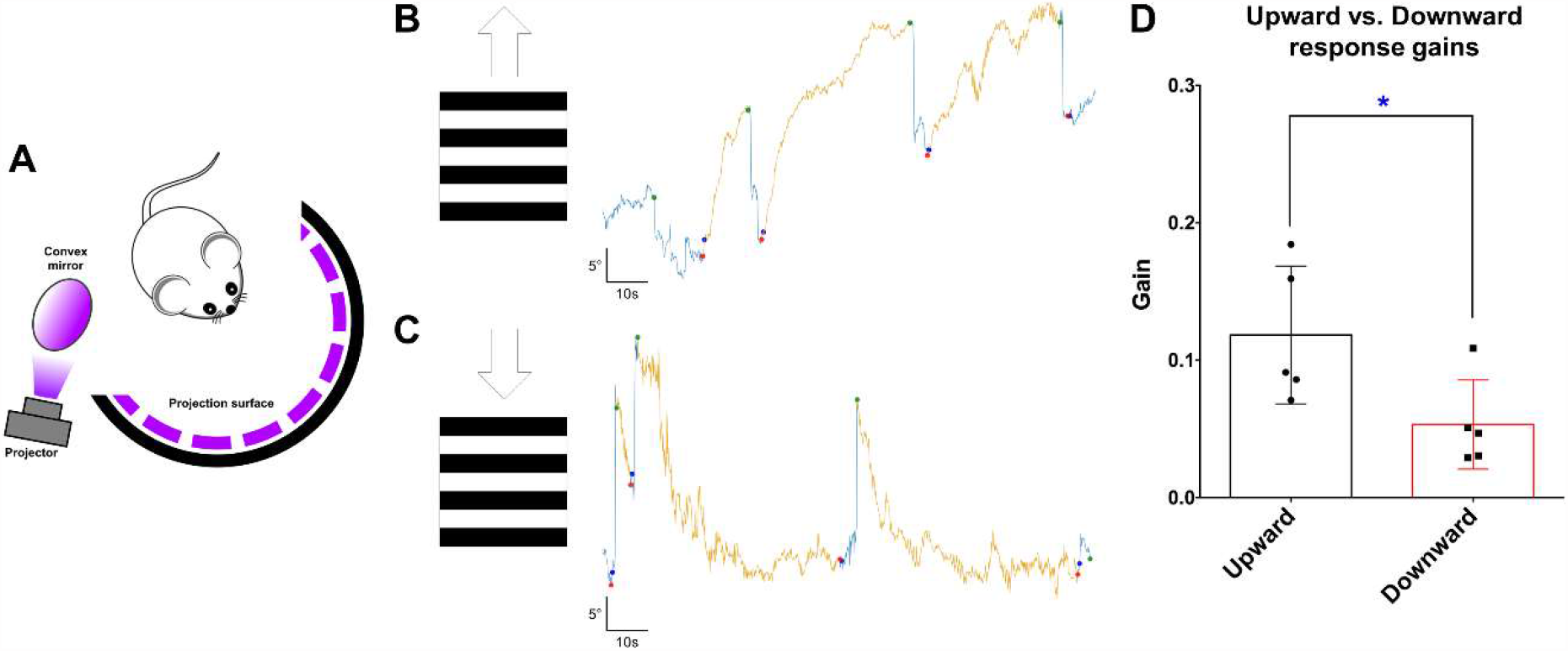
Application of analysis program to alternative video-oculography methods. **A)** Set-up of virtual drum for OKR stimulation as described^10^. A 405nm wavelength DLP projector was reflected via a convex mirror onto a hemisphere to create a virtual drum. Unidirectional bar gratings are shown to a head-fixed animal in vertical directions. **B-C)** Upward (**B)** and downward (**C**) tracking phases are identified and selected for quantitative analysis. Slow phases are highlighted in yellow. **D)** Tracking gains calculated from vertical tracking of wildtype animals (n=5) using described method. Asymmetric tracking ability is observed with a significant decrease of downward tracking. Data are presented as mean ± SD. Data analyzed with an unpaired T-test. *p<0.05.

## DISCUSSION

The method of OKR quantification presented here provides several novel advantages for studying murine visual responses reflected in eye tracking. Direct assessment of eye tracking gains provides a more accurate characterization of eye movement than traditional manual counting of fast phase saccades, and our algorithm provides a faster, more efficient, and reproducible quantification method than previous gain analyses. Although useful, the counting of saccades provides an indirect assessment of the actual tracking motion whereas our method provides direct quantification of tracking speeds, which takes into account additional parameters including saccade amplitudes and the length of slow phases. In addition, fast phase saccade counting relies on the assessment of saccade frequency over a set period of time, requiring that each trace has little to no noise, such as random saccades from animal stress. This results many collected traces being unusable and therefore adds significant time for data collection in order to accurately assess eye tracking. However, our method addresses this issue by directly quantifying individual slow phases within a given trace. Since saccadic frequency is no longer the defining parameter for analysis, traces with some level of noise can still be analyzed and will be accurately quantified. This allows for more usable collected data and thus higher statistical power for the data collected, increasing accuracy and reproducibility when quantifying eye tracking and characterizing OKR parameters for individual mice and different groups of mice.

In addition, our standardized, accessible, and semi-automated analysis pipeline facilitates higher-throughput and reproducible quantification, as compared to manual ETM counting. Even using previous methods, sorting through OKR traces is time-consuming and labor-intensive. Our method allows not only for faster trace quantification, but it also reveals additional OKR parameters, with direct quantification of the distance the eye has traveled, its velocity, and its speed relative to the stimulus in both horizontal and vertical components of the movement vector. Further, given the automated identification of saccades and calculation of tracking speeds afforded by our method, potential experimenter bias in data analysis is greatly reduced compared to other quantification methods such as manual ETM counting.

Finally, our method is adaptable to other video-oculography collection methods and provides a robust analysis platform that can be easily tailored to a laboratory’s specific needs. Through analysis of wildtype and existent mutant OKR data, and also wildtype OKR data collected using different methods, we show that our analysis tool is capable of: a) quantifying OKR tracking ability with high accuracy and speed; b) identifying behavioral differences resulting from genetic perturbations of the visual system; and c) assessing visual wave data from different video-oculography methods. The accessibility and general adaptability of our analysis platform facilitates further study OKR responses and will enhance behavioral studies that characterize neural circuit assembly and dynamics in the context of oculomotor responses.

To obtain accurate measurements and subsequent useful data analyses several steps are necessary for data collection. We recommend collecting OKR data in multiple sessions to allow for the animal to acclimate to the behavioral testing apparatus, reducing the impact of animal stress on behavioral responses. During OKR data collection, proper calibration of the video-oculography equipment is critical for accurate quantification, since analyzed data is a direct function of the processed wave. Additionally, the use of the Spyder IDE is necessary for graph supervision through matplotlib. One limitation of our method is that it primarily assesses behavioral responses to unidirectional gratings or bars in the four cardinal directions. Current work is underway to incorporate analysis capabilities for additional stimuli, including oscillatory sinusoidal stimuli. However, given the accessibility and framework design of our platform, the tools are available for others to expand its capabilities and tailor this platform for distinct behavioral experimental paradigms.

In conclusion, we describe here a new, accessible, and versatile tool for analysis of mouse OKR behavioral responses in more depth and with more quantitative power than is available with existing methodologies. This method can be easily used by novice Python users and contains an established analysis pipeline and interface to analyze OKR waves rapidly and accurately with increased rigor and reproducibility. Additionally, the adaptability of the method provides an adaptive framework that users can easily tailor to fit their specific needs and data collection set-ups. We anticipate that this quantitative method will advance study of murine oculomotor responses and help in the understanding the development and function of neural circuitry that drives visual system behaviors.

## ACKNOWLEDGMENTS

This work was supported by R01 EY032095 (ALK) and a VSTP pre-doctoral fellowship 5T32 EY7143-27 (JK).

